# Functional and genomic adaptations of blood monocytes to pregravid obesity during pregnancy

**DOI:** 10.1101/2020.12.02.409185

**Authors:** Suhas Sureshchandra, Nicole E. Marshall, Norma Mendoza, Allen Jankeel, Michael Z. Zulu, Ilhem Messaoudi

**Affiliations:** Department of Molecular Biology and Biochemistry, University of California, Irvine CA 92697; Institute for Immunology, University of California, Irvine CA 92697; Center for Virus Research, University of California, Irvine CA 92697; Maternal-Fetal Medicine, Oregon Health and Science University, Portland OR 97239

**Keywords:** Pregnancy, obesity, monocytes, inflammation, chromatin, gestation, transcription, epigenome, tolerance

## Abstract

Pre-pregnancy obesity is associated with several adverse maternal health outcomes, notably increased risk of infection as well as the incidence of gestational diabetes, preeclampsia, and preterm birth. However, the mechanisms by which pregravid obesity disrupts the pregnancy associated “immune clock” are still unknown. To address this question, we collected blood samples from women during the first and third trimesters and determined the impact of both pregnancy and pregravid obesity on circulating immune mediators, immune cell subset frequencies, and peripheral immune responses. While regardless of BMI, pregnancy was associated with an elevation in both Th1 and Th2 cytokines, pregravid obesity was associated with a dysregulation in circulating myeloid factors at term. Moreover, pregnancy in lean subjects was associated with enhanced monocyte activation, augmented chromatin accessibility at inflammatory loci, and heightened responses to LPS. Pregravid obesity disrupted this trajectory and was accompanied by a lack of transcriptional and epigenetic changes and alterations in metabolic status strongly suggesting a skewing towards immunotolerance. These findings provide novel insight into the increased susceptibility to infections observed with obesity during pregnancy.

**SUMMARY:** A healthy pregnancy is associated with progressive innate immune activation. Maternal factors such as obesity compromise this myeloid cell activation trajectory at genomic, epigenomic, functional, and metabolic levels, resulting in stagnant immune responses, suggestive of a state of tolerance.

## INTRODUCTION

The maternal immune system undergoes several fine-tuned adaptations over the course of pregnancy (Mor and Cardenas, 2010). These changes are believed to facilitate implantation, fetal tolerance, fetal growth and development, and finally labor and parturition without compromising protection against microbial infections (Mor et al., 2011). A recently described “immunological clock” of pregnancy suggests that peripheral immune adaptations do not necessarily follow waves of pro- and anti-inflammatory phases (Mor and Cardenas, 2010), but rather a progressive activation of signaling molecules (Aghaeepour et al., 2017) and alterations in the circulating proteome (Aghaeepour et al., 2018). Specifically, while the peripheral adaptive immune system is skewed towards Th2/Treg responses and away from Th1/Th17 responses (Luppi et al., 2002b; Polese et al., 2014; Sacks et al., 2003), innate immune cells are progressively activated over the course of a healthy pregnancy (Aghaeepour et al., 2017). This activation is manifested through higher production of pro-inflammatory cytokines IL-6, IL-1β, and IL-12 by peripheral blood mononuclear cells (PBMC) following *ex vivo* stimulation with LPS and bacteria (Aghaeepour et al., 2017; Faas et al., 2014; Luppi et al., 2002a; Naccasha et al., 2001; Sacks et al., 2003; Sacks et al., 1998) or viral particles (Kay et al., 2014; Le Gars et al., 2016). However, the precise mechanisms underlying this progressive activation of the innate immune branch remain poorly defined.

A third of women of reproductive age in the US meet the definition of obese (body mass index (BMI)>30) (Flegal et al., 2016), making pregravid obesity one of the most critical and common health challenges during pregnancy. High pregravid BMI increases the risk for gestational diabetes (Chu et al., 2007; Torloni et al., 2009), gestational hypertension, preeclampsia (Huda et al., 2014; Wang et al., 2013), early pregnancy loss, placental abruption, abnormal fetal growth, premature labor, and stillbirth (Aune et al., 2014; Kim et al., 2016). High pregravid BMI is also an independent risk for infection during pregnancy (urinary and genital tract infections, sepsis) (Sebire et al., 2001), labor (chorioamnionitis) (Cnattingius et al., 2013; Hadley et al., 2019), and post-partum (surgical site infections following cesarean) (Conner et al., 2014; McLean et al., 2012).

These adverse outcomes suggest a dysregulated immune status in pregnant women with obesity; but the impact of pregravid obesity on the pregnancy “immune clock” remains poorly defined. Pregravid obesity is marked by elevated circulating levels of inflammatory mediators C reactive protein (CRP) and IL-6 (Basu et al., 2011a; Sureshchandra et al., 2018), subclinical endotoxemia, and signs of heightened inflammation in the adipose tissue compartment (Basu et al., 2011a). Increased exposure to subclinical levels of LPS with obesity could potentially alter both phenotype and fitness of innate immune cells. Indeed, we recently reported dysregulation in secretion of immune mediators, especially those expressed by monocytes and dendritic cells (DCs), following *ex vivo* stimulation with TLR agonists of PBMC collected at gestational week 37 from subjects with obesity (Sureshchandra et al., 2018).

Amongst innate immune cells, monocytes and macrophages play critical roles at various stages of pregnancy. Circulating monocytes play seminal roles in implantation, placentation, and labor, while decidual monocyte-derived macrophages play critical roles in angiogenesis, tissue development, and wound healing at the maternal fetal interface (Basu et al., 2011b; Mor and Abrahams, 2003; Mor et al., 2011). Monocytes/macrophages have also been implicated in several obesity-associated complications during pregnancy such as preeclampsia (Faas et al., 2014), preterm labor (Gomez-Lopez et al., 2014), poor placental development (Faas and de Vos, 2017), and chorioamnionitis (Ben Amara et al., 2013). However, several questions still remain unaddressed: (1) how does pregravid obesity alter monocyte responses and function throughout pregnancy; (2) what are the cellular and molecular mechanisms of underlying those changes?

To answer these questions, in this study, we defined longitudinal changes in circulating immune mediators, peripheral innate immune cell subsets and responses of monocytes and DCs to *ex vivo* stimulation during early and late stages of gestation in both lean and obese women to elucidate the impact of pregravid obesity on the “pregnancy immunological clock” using a combination of immunological, transcriptional and epigenetic analyses. We report a gestation-associated trajectory of monocyte activation that is accompanied by an enhanced response to LPS and poising of the epigenome towards a state of heightened activation. Pregravid obesity disrupts this trajectory of monocytes’ activation, skewing them towards an immunoregulatory phenotype characterized by attenuated responses to LPS and lack of epigenetic and metabolic plasticity observed in lean pregnancy. This dysregulation in monocyte activation with pregravid obesity was accompanied with systemic changes in circulating cytokine and chemokine environment consistent with a regulatory environment. Collectively, these findings provide novel insight into mechanisms by which pregravid obesity disrupt pregnancy associated adaptations that mediate tolerance while protecting the developing fetus and pregnant woman against microbial challenges.

## RESULTS

### Pregnancy and pregravid obesity alter the circulating metabolic and inflammatory environment

Both pre-pregnancy BMI and fat mass were recorded to stratify subjects as lean (BMI < 25) and obese (BMI >30). Patient demographics are summarized in Table 1. Pregravid BMI correlated strongly with fat mass percentages, both at T1 and T3 (Supp Figure 1A). Gestational weight gain was significantly lower in the obese group (Supp Figure 1B). We measured plasma levels of circulating metabolic hormones, adipokines, cytokines, chemokines, and growth factors at T1 and T3 (Figure 1A). At both T1 and T3, high pregravid BMI was associated with elevated circulating CRP (Figure 1B) and IL-6 (Figure 1C) levels. Systemic levels of insulin (Figure 1D) and leptin (Figure 1E) were significantly higher in the obese group at both time points. Interestingly, their levels remained unchanged in the obese group but increased with gestation in the lean group (Figures 1D and 1E). Plasma levels of resistin were also higher in the obese group at both time points but did not increase with pregnancy in either group (Supp Figure 1C). Finally, plasma levels of the adipokines adipsin (Supp Figure 1D) and adiponectin (Supp Figure 1E) increased with pregnancy in both groups and no differences between the groups were noted. No differences were detected in plasma levels of gut hormone peptide YY (PYY) with either pregnancy or pregravid obesity (Supp Figure 1F).

**Figure 1:**
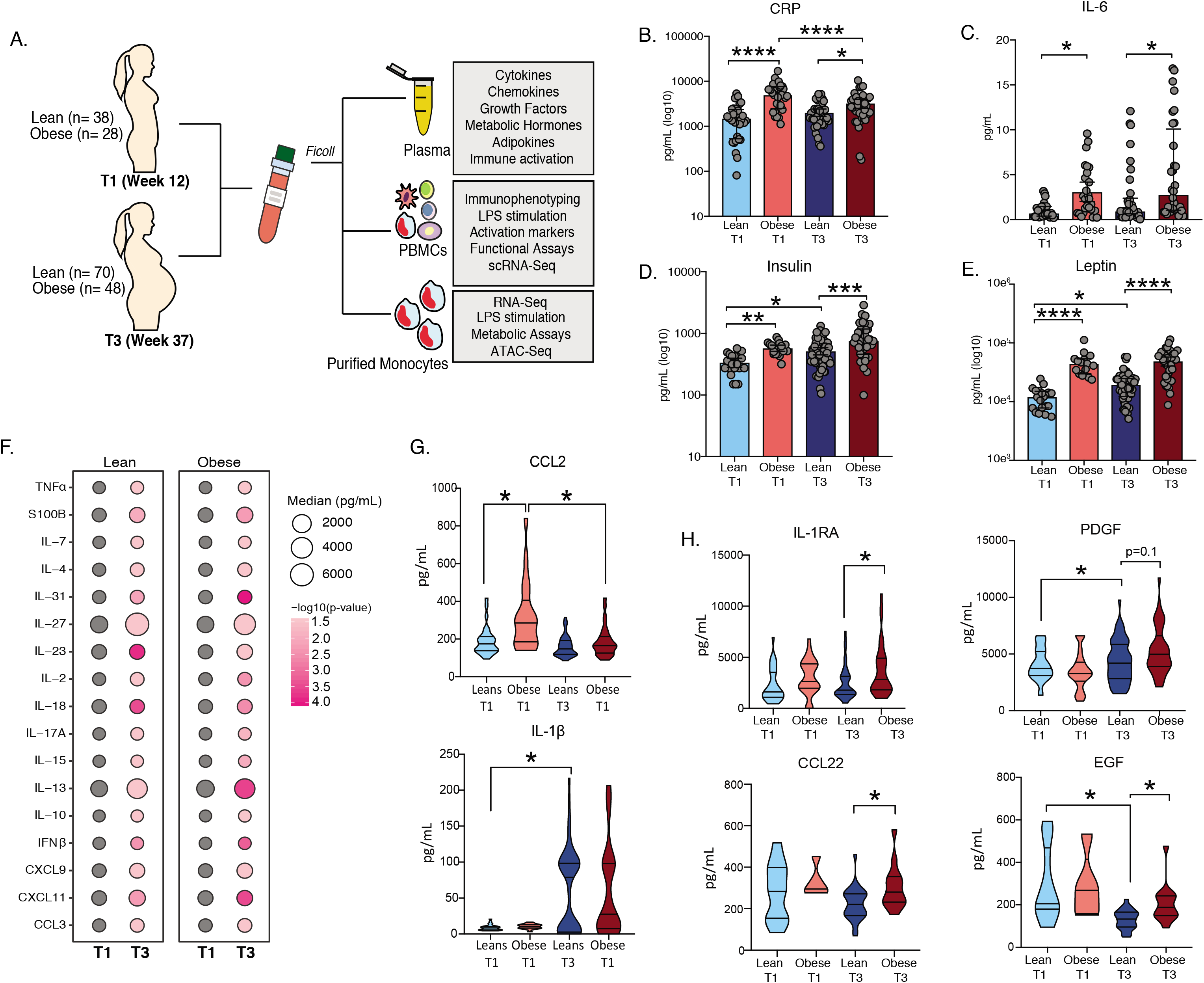
Experimental Design and longitudinal changes in maternal inflammatory environment. (A) Blood samples were obtained from pregnant women during early (week 12 — T1) and late (week 37 — T3) pregnancy. PBMC and plasma were isolated and used to assess the impact of pregravid obesity on maternal immunity. (B, C) Bar graphs depicting changes in plasma levels of inflammatory mediators (B) CRP and (C) IL-6 measured using high sensitivity ELISA. (D, E) Circulating levels of metabolic hormones (D) insulin, (E) leptin were measured using luminex. Levels of significance: * – p<0.05, ** – p<0.01, *** p<0.001, **** – p< 0.0001. Bars denote median with interquartile ranges. (F) Bubble plot representing changes in immune mediator levels in plasma with gestation. Size of the bubble represents median values of analytes in pg/mL (log2 transformed). The colors at T3 represent the levels of significance relative to T1. (G-H) Four-way violin plots comparing plasma levels of (H) pro-inflammatory myeloid factors CCL2 and IL-1β and (H) regulatory cytokine (IL-1RA), chemokine (CCL22), and growth factors (PDGF and EGF). Levels of significance: *– p<0.05, ** – p<0.01; *** – p<0.001; **** – p<0.0001.

**Table 1:** Cohort characteristics

|  | **Lean (n=69)** | **Obese (n=48)** | **P-Value** |
| --- | --- | --- | --- |
| **Age (y)** | 32.8±4.7 | 29.4±9 | n.s |
| **Race:** American<br>Indian/Alaskan Native | 2 | 4 |  |
| **Race:** Asian American | 6 | 1 |  |
| **Race:** African American | 0 | 6 |  |
| **Race:** Pacific Islander | 2 | 0 |  |
| **Race:** White/Caucasian | 64 | 37 |  |
| **Ethnicity:** Hispanic | 6 | 8 |  |
| **BMI (kg/m<sup>2</sup>)** | 21.9±1.7 | 35.2±5.2 | <0.0001 |
| **Fat mass at 12 weeks**<br>**(kg)** | 17.3±4.1 | 42.6±11.4 | <0.0001 |
| **Fat mass at 37 weeks**<br>**(kg)** | 22±5.5 | 44.2±11.1 | <0.0001 |
| **GWG (kg)** | 15±4.5 | 10.4±7.6 | <0.001 |

Independent of maternal BMI, pregnancy was associated with shifts in the circulating inflammatory environment from T1 to T3 (Figure 1F). Levels of both pro-inflammatory (TNFα, IL-2, IFNβ, IL-7, IL-17A, IL-18, IL-23) and regulatory (IL-4, IL-10, and IL-13) cytokines, as well as chemokines (CCL3, CXCL9, and CXCL11) significantly increased during pregnancy in both lean and obese groups (Figure 1F). At T1, obesity was associated with elevated levels of myeloid cell chemoattractants CCL2 (Figure 1G) and CCL4. Levels of CCL2 decreased dramatically in the obese group, but remained unchanged in the lean group, with pregnancy (Figure 1G). In contrast, levels of CCL4 as well as the pro-inflammatory myeloid cell activator IL-27 increased with gestation in both groups, albeit more prominently in the lean group (Supp Figure 1H, Supp Figure 1G). Levels of the myeloid pro-inflammatory factor IL-1 β only increased in the lean group with pregnancy (Figure 1F). Additionally, levels of regulatory factors IL-1RA, chemokine CCL22, and growth factors PDGF and EGF were increased at T3 only in the obese group (Figure 1H). Finally, a significant drop in regulatory factor VEGF with pregnancy was observed in lean group alone (Supp Figure 1I).

We next correlated pregnancy-associated changes in circulating factors with maternal BMI (Supp Figure 1J). Our analyses revealed positive association between maternal BMI and changes in inflammatory factors leptin and IL-18 as well as regulatory factors IL-10 and PDGF (Supp Figure 1J). Surprisingly, though CRP levels remained elevated with obesity at both T1 and T3 (Figure 1B), the decrease negatively correlated with maternal BMI (Supp Figure 1J).

### Pregravid obesity alters the frequencies of monocyte subsets and disrupts pregnancy-associated activation trajectory of circulating monocytes

Since some of the most interesting differences between the lean and obese groups with obesity were myeloid factors (CCL4, IL-1β, IL-1RA, IL-27), we next investigated if pregravid obesity altered frequency of circulating innate immune cell subsets. Complete blood cell (CBC) counts revealed increased numbers of white blood cells (WBC) with pregnancy and with pregravid obesity (Supp Figure 2A). Differences in WBC numbers with obesity were primarily driven by granulocytes, with no major differences in lymphocyte or monocyte numbers (Supp Figure 2A).

Multi-parameter flow cytometry analysis of innate immune cells in PBMC (Supp Figure 2B) revealed no changes in major cell populations (Supp Figures 2C-2E) with pregnancy or pregravid obesity with the exception of a decrease in dendritic cells (DC) frequency in the lean group (Supp Figure 2D). In mothers with obesity, pregnancy was associated redistribution of DC subsets, with higher mDC (Supp Figure 2F) but lower pDC subsets at T3 (Supp Figure 2G). Pregravid obesity was also associated with a reduced proportion of nonclassical (CD16+) subset of monocytes (Figure 2A). Although the frequency of NK cells was unchanged with pregnancy or pregravid obesity (Supp Figure 2E), the relative abundance of CD56^bright^CD16^+^ NK cells increased with pregnancy in both groups (Supp Figure 2H). Very few changes in the frequency of lymphocytes were observed with pregnancy and/or obesity (Supp Figure 3). Pregravid obesity was associated with a slight decrease in the frequency of B cells (Supp Figure 3A), that was largely mediated by a drop in the frequency of memory and marginalzone (MZ) like B cells (Supp Figure 3B). Pregnancy in the lean group was also accompanied by a decrease in naïve and MZ like B cells (Supp Figure 3B) as well as transitional effector memory CD4+ T cells (Supp Figure 3E).

**Figure 2:**
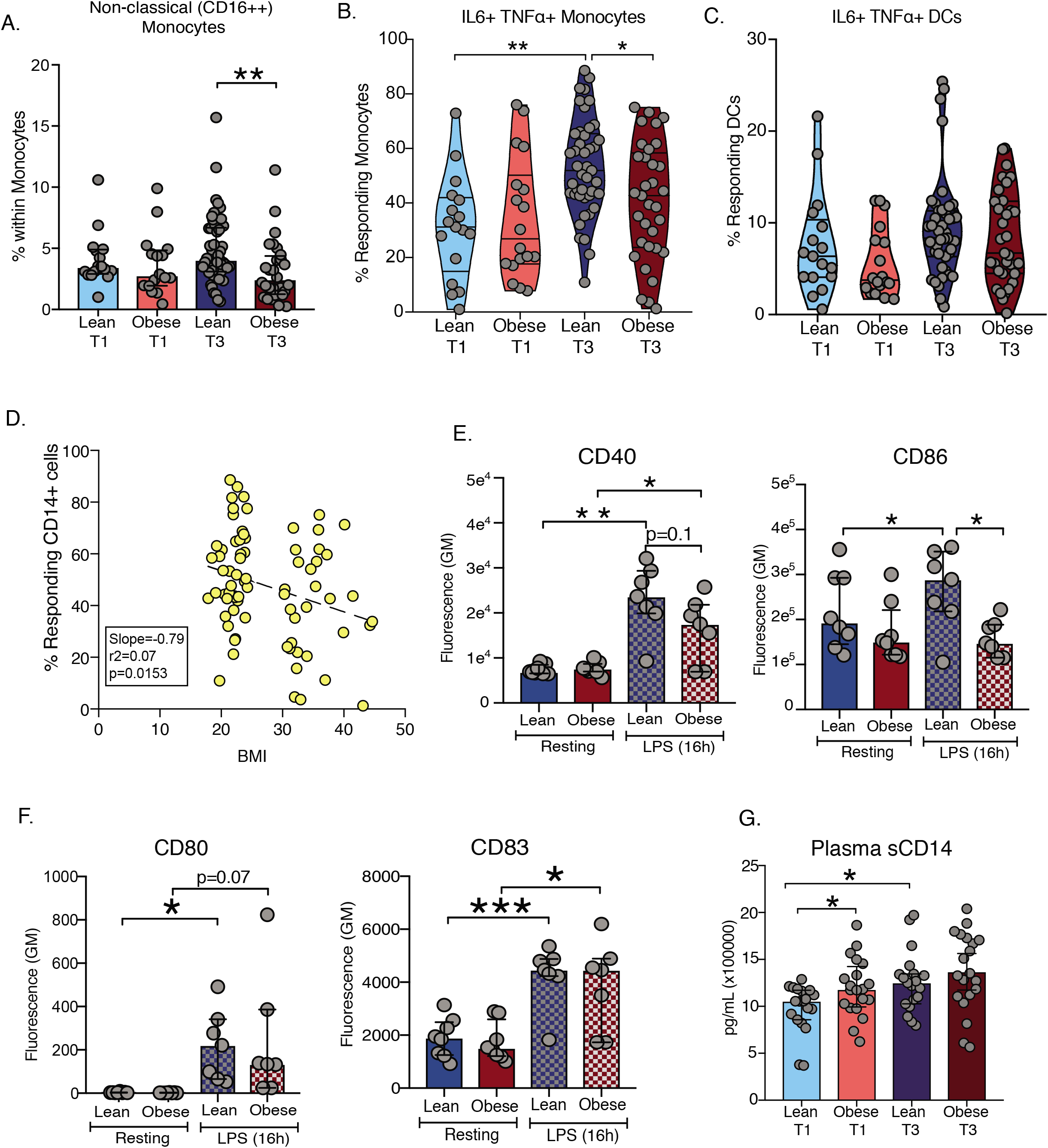
Pregnancy and obesity associated changes in innate immune phenotype and *ex vivo* response. (A) Bar graph depicting percentages of non-classical subset (CD14^+^CD16^++^) within monocytes. (B-C) Violin plots showing percentage of IL6 and TNF producing (B) monocytes and (C) dendritic cells (DCs) following 16h LPS stimulation. (D) Linear regression analysis of maternal BMI and monocyte responses to LPS at T3. Deviations in slope of regression line were tested using F-test. (E-F) Surface expression of activation markers (E) CD40, CD86 and (F) CD80, and CD83 on monocytes following 16h LPS stimulation at T3 time point. (G) Plasma levels of soluble CD14 measured at both time points using ELISA. Levels of significance: * – p<0.05, ** – p<0.01. Bars denote median and interquartile ranges.

We next investigated the impact of pregnancy and pregravid obesity on the ability of innate immune cells and T cells to respond to *ex vivo* stimulation with lipopolysaccharide (LPS) and anti-CD3/CD28 respectively (Figure 1A) using intracellular staining (Supp Figure 4A). Pregnancy in lean women was associated with increased frequency of IL6^+^TNF^+^ producing monocytes (Figure 2B), but not dendritic cells (Figure 2C), in response to LPS stimulation. This enhanced monocyte response was absent in the obese group resulting in fewer responding monocytes compared to the lean group at T3 (Figure 2C). Indeed, frequency of responding monocytes at T3 negatively correlated with maternal pregravid BMI (Figure 2D). Enhanced response to LPS at T3 by monocytes from the lean group was accompanied by a higher induction of activation markers CD40 and CD86 (Figure 2E) as well as increased levels of circulating soluble CD14 (sCD14), a surrogate for *in vivo* monocyte activation (Figure 2G). Activation markers CD80 and CD83 were upregulated to similar levels in both groups (Figure 2F). Cytokine production by T cells remained largely unchanged with pregnancy and pregravid obesity (Supp Figures 4) with the exception of enhanced Th2 responses (Supp Figure 4B).

### Single cell RNA-seq reveals significant shifts in monocyte phenotypes at T3

To better understand the impact of pregravid obesity on monocyte activation with pregnancy, we first carried out bulk RNA Seq analysis on purified monocytes obtained from T1 and T3 from lean and obese subjects (n=3/time point). Pairwise longitudinal comparison of the transcriptome of resting monocytes from the lean group showed modest changes, with increased expression of MHC molecules (*B2M, HLA-DRA*) and proinflammatory genes (e.g. *VCAN, LYZ*) (P-value <0.001) at T3 relative to T1 (Supp Figure 5A). On the other hand, monocytes from mothers with obesity (n=3/time point) exhibited modest suppression of genes involved in of interferon signaling (*STAT1, GBP5, PSMB8*) and response to reactive oxygen species (*STRBP, HEBP2, TRAP1, TRPA1*) (P-value <0.001) with gestational age (Supp Figure 5B).

Given the modest shifts detected using bulk RNA Seq and the significant functional differences observed at T3, we performed single cell RNA-sequencing of total PBMC (n=2/group) obtained at delivery (T3). After concatenating all 4 libraries, we computationally extracted the monocyte clusters (~600 cells/group – total 2400 cells) using expression of *CD14* (clusters 7, 11, and 16 in the initial UMAP of PBMC; Supp Figure 5C). Following subsequent iterations of doublet removal and UMAP clustering, we identified 6 major clusters of monocytes (Figure 3A) representing the three subsets traditionally identified by flow cytometry based on expression of *CD14, FCGRA* (CD16), and *HLA-DRA* (Supp Figure 5D): classical monocytes (CM – clusters I, III, V); nonclassical monocytes (NCM – clusters II and IV); and intermediate (IM-cluster VI) (Figure 3B). Trajectory analysis using Monocle confirmed this classification placing non-classical monocytes at the end of the pseudotime with intermediate monocytes transitioning from classical to non-classical subsets (Figure 3C and Supp Figure 5E). Pregravid obesity was associated with a dramatic drop in the number of non-classical monocytes (especially cluster IV) (Figures 3D and 3E). We also detected a significant increase in classical monocytes mediated by a substantial increase in cluster III but a decrease in cluster V (Figures 3D and 3E). Classical monocytes within cluster III expressed markers of insulin resistance (*CD44, LMNA, THBD*) (Figure 3B), while those within cluster V (Figures 3B, 3D and 3E) expressed pro-inflammatory genes (*VCAN, S100A9, LYZ*) (Figure 3B).

**Figure 3:**
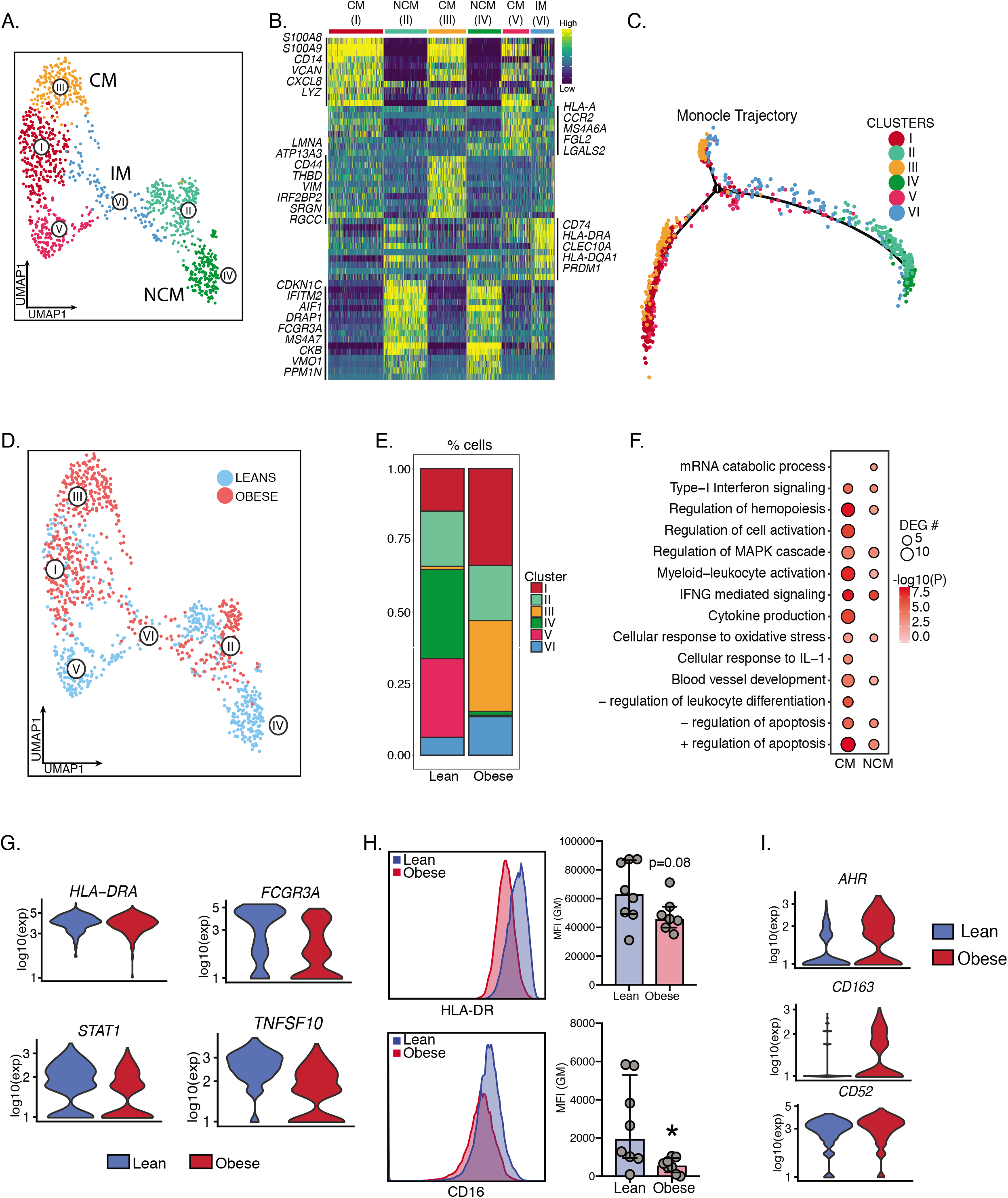
Single cell profiles of blood monocytes at delivery. (A) UMAP of monocytes following extractions of CD14^high^ clusters from UMAP of PBMC from lean (n=2) and obese (n=2) subjects, colored by cell state (clusters), and annotated by cluster numbers and monocyte subsets (CM-classical monocytes, IM – intermediate, NCM – nonclassical monocytes). (B) Heatmap showing expression patterns of the 10 most distinguishing markers per cell state with selected markers indicated (yellow = high expression; green = intermediate expression; blue = low expression) phenotype. (C) Trajectory analysis of monocyte with cells colored by their original cluster designation. (D) UMAP visualization of monocytes, colored by group. (E) Relative frequencies of cells within each UMAP cluster in either group. (F) Functional enrichment of genes differentially expressed with pregravid obesity in classical and non-classical monocyte subsets. Bubble size and color are indicative of number of genes within each ontology term and the significance of its enrichment. (G) Violin plots showing log-transformed, normalized expression levels for select genes down-regulated with pregravid obesity (H) Surface expression of HLA-DR and CD16 using flow cytometry. Error bars represent medians with interquartile range. (I) Violin plots showing log-transformed, normalized expression levels for select genes up-regulated with pregravid obesity

Differential gene expression analysis within both classical and non-classical clusters revealed transcriptional changes associated with immune activation, signaling, differentiation, and apoptosis in with obesity (Figure 3F). Specifically, we detected suppression of *HLA-DRA, STAT1, TNFSF10,* and *FCGR3A* (Figure 3G) with pregravid obesity. Reduced surface expression HLA-DR and CD16 (encoded by *FCGR3A*) with pregravid obesity was confirmed using flow cytometry (Figure 3H). As observed with bulk RNA sequencing (Supp Figure 5B), we observed a global suppression of interferon signaling pathways with pregravid obesity across all clusters (Supp Figure 5F) as exemplified by reduced expression of interferon-stimulated genes (*IFIT3, ISG15, OAS1*) (Supp Figure 5G). On the other hand, pregravid obesity was associated with up-regulation of immuno-regulatory molecules such as *AHR, CD52,* and *CD163* (Figure 3I).

### Defects in monocyte responses with maternal obesity at T3 are cell intrinsic

We next investigated whether reduction in cytokine production by circulating monocytes in response to LPS with pregravid obesity is cell intrinsic. Monocytes were isolated from PBMC (Supp Figure 6A) obtained from lean subjects and those with obesity at T1 and T3 and stimulated with LPS for 8 hours (Figure 4A). Significant differences in the monocyte response to LPS with pregravid obesity were detected at both T1 (Supp Figure 6B) and T3 (Supp Figure 6C). Although gene expression responses to LPS were more robust at the T3 time point compared to T1 for both the lean (Figure 4B) and obese groups (Figure 4C), DEG detected at T3 in the obese group were predominantly down regulated (Figure 4C). Moreover, functional enrichment revealed a higher proportion of DEG detected at T3 in the lean group enriched to GO terms associated with myeloid activation, leukocyte adhesion, cytokine signaling (Figure 4D). In line with these observations, we detected lower induction of key inflammatory molecules *IL6, IL1B, TNF,* and *CCL3* (Figure 4E) in the obese group. Furthermore, at T3, we observed reduced expression of key transcription factors *NFKB1, FOSB, NCOR, CIITA* (Figure 4F) with pregravid obesity. These findings were supported by transcription factor analysis using ChEA3, which revealed reduced number of RELA, IRF, and STAT-regulated DEG with pregravid obesity, particularly at T3 (Figure 4G). On the other hand, the number of LPS-responsive genes regulated by TF IRF1, STAT1, and STAT3 were increased in the lean group at T3 compared to T1 (Figure 4G). Finally, we report poor induction of key regulators of chromatin reorganization (*KDM2B, SIRT1, SIRT2, EP300, KDM4C*) following LPS stimulation in obese group at T3 (Figure 4D and 4H), suggestive of epigenetic constraints to immune activation.

**Figure 4:**
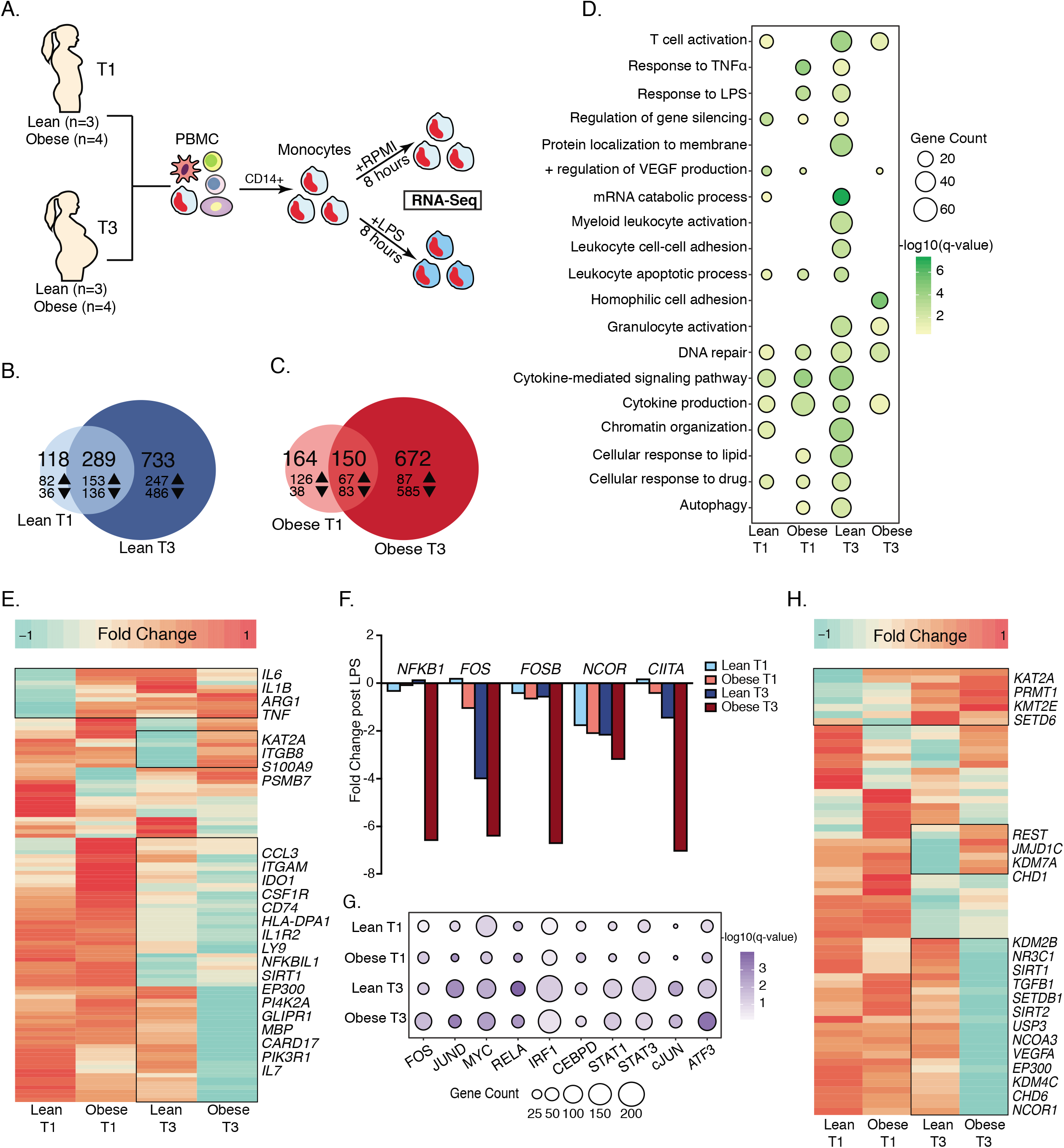
Cell intrinsic defects in monocyte responses to LPS with maternal obesity. (A) Experimental design for RNA-seq. (B-C) Venn diagrams comparing LPS induced differentially expressed genes (DEG) between T1 and T3 in purified monocytes from (B) lean and (C) obese groups following 8h LPS exposure. Numbers of Up- and down-regulated DEG (FDR≤0.05) are annotated with corresponding arrows (D) Bubble plot representing gene ontology terms enriched following LPS stimulation in all four groups following LPS stimulation. Both up- and down-regulated genes are included. Size of the bubble represents numbers of genes mapping to the term while color indicates level of significance. (E) Heatmap comparing fold changes of DEG (both up- and down-regulated) that mapped to GO terms “cytokine production” and “Myeloid leukocyte activation”. Green indicates down-regulation while orange indicates up-regulation. (F) Fold change of key nuclear factors significantly down-regulated with LPS. (G) Bubble plots representing number of DEG regulated by specific LPS inducible transcription factors predicted by ChEA3. Size of the bubble represents the numbers of DEG regulated by each transcription factor. Color of the bubble represents the level of significance for each prediction. (H) Heatmap comparing fold changes of DEG (both up- and down-regulated) that mapped to GO terms “Chromatin organization”. Green indicates down-regulation while orange indicates up-regulation.

### Pregravid obesity disrupts pregnancy-associated epigenetic trajectory of circulating monocytes

We next asked if there exists a pregnancy-associated epigenetic clock that explains the enhanced monocyte responses to LPS at T3 relative to T1 in lean subjects. We therefore profiled open chromatin of purified resting monocytes isolated from lean and obese group at both time points using ATAC-Seq (Figure 1A). Principal component analysis revealed significant shift in ATAC profiles of monocytes from T1 to T3 in lean subjects (Figure 5A), with 7132 regions open at T3 relative to T1 (Figure 5B), most of which overlapping promoter regions (Figure 5C). Promoter regions open in leans at T3 relative to T1 regulated genes involved in leukocyte activation, myeloid differentiation, and regulation of oxidative stress (Supp Figure 7A). Furthermore, open intergenic regions in leans at T3 relative to T1 regulated genes involved in mounting inflammatory response to lipids and LPS (Supp Figure 7B).

**Figure 5:**
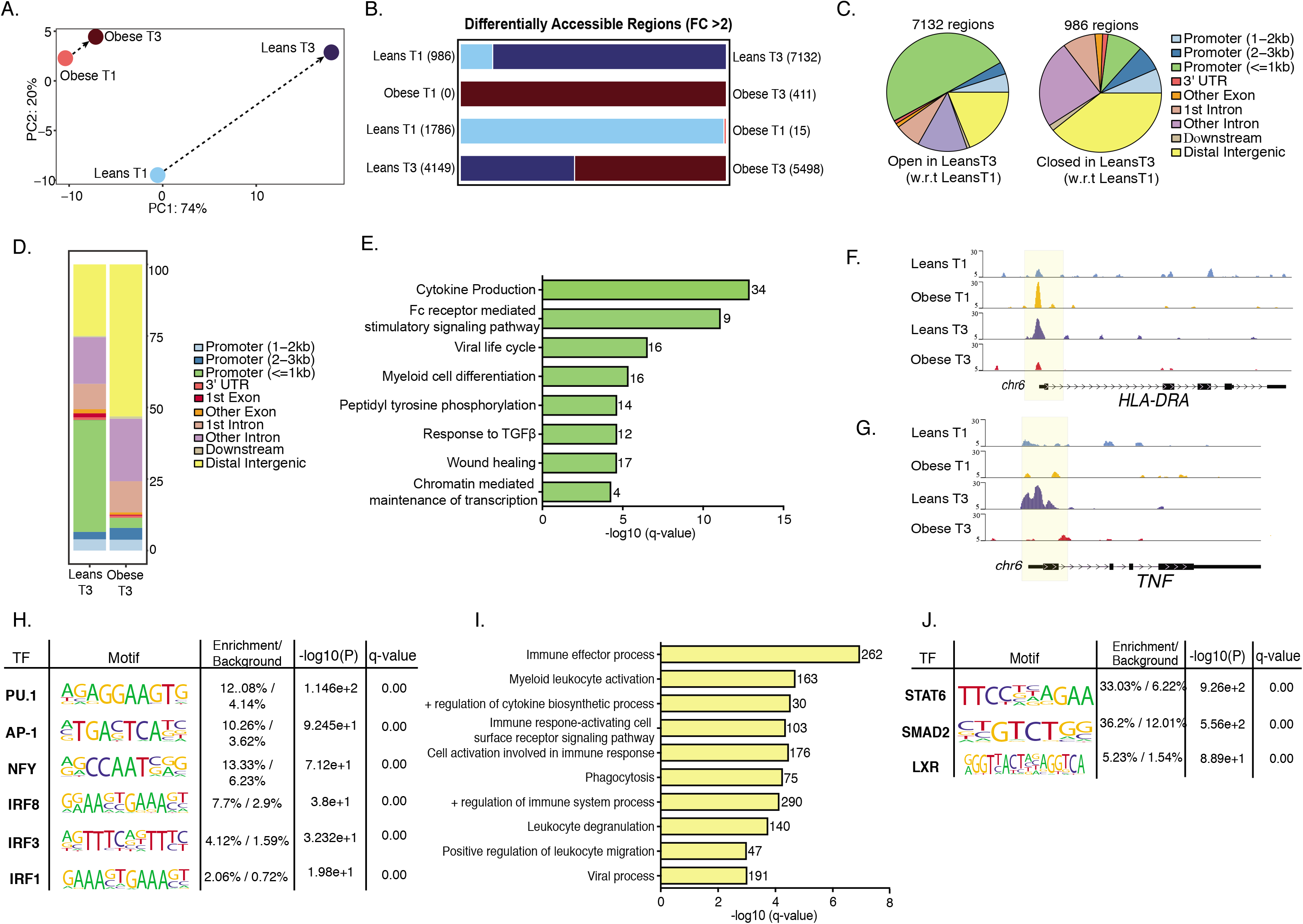
Epigenetic adaptations with pregnancy and maternal obesity. (A) PCA of chromatin accessibility profiles in monocytes isolated at T1 and T3 from lean and obese subjects. Arrows represent trajectory from T1 to T3. (B) Bar graphs with numbers of differentially accessible regions (DAR) identified in each comparison. Numbers in parenthesis denote open regions identified in a particular group relative to the comparison group. (C) Pie charts showing genomic contexts of loci identified as DAR with gestation in the lean group. (D) Genomic annotations of 4149 and 5,498 DAR identified as open in lean and obese group respectively at T3. (E) Functional enrichment of genes regulated by promoter-associated DAR open in lean group relative to obese group (FC>2). Genes associated with these regions are quantified next to each term (F-G) WashU Epigenome tracks for (F) *HLA-DRA* and (G) *TNF* locus with promoter vicinity highlighted in yellow. (H) Enrichment of motifs in promoter associated open DAR in lean group relative to the obese group at T3. Only motifs identified in myeloid cells are included in the table. (I) Gene ontologies of intergenic associations identified by GREAT as significantly open in lean group relative to obese group at T3. (J) Enrichment of motifs in open intergenic DAR in obese group relative to lean group at T3. Only motifs identified in myeloid cells are included in the table.

In contrast, limited changes in chromatin accessibility were noted between T1 and T3 in the obese group (Figures 5A and 5B). Direct comparisons of lean and obese groups at each time point revealed more dramatic differences in chromatin accessibility at T3 (Figure 5D) than at T1 (Supp Figure 7C). Specifically, at T3, 40% of loci open in lean group relative to the obese group overlapped promoter regions of genes involved in cytokine production, Fc receptor mediated signaling, and wound healing (Figure 5E). This is further illustrated by pileups of the loci of inflammatory genes *HLA-DRA* (Figure 5F), *TNF* (Figure 5G), *IL6ST* (Supp Figure 7D), and activation marker *CD80* (Supp Figure 7E). Finally, motif analysis revealed that promoter regions open in monocytes from lean subjects at T3 (closed in monocytes from subjects with obesity) harbor binding sites for transcription factors PU.1 and AP-1 and interferon regulators IRF1, IRF3, and IRF8. The limited number of promoter-associated regions that were open in obese group relative to leans at T3 (Figure 5D) regulated genes primarily involved in histone methylation, and metabolic processes (Supp Figures 7F and 7G).

GREAT enrichment of intergenic loci closed in obese group at T3 relative to leans (Figure 5H) revealed several associations with immune effector process, phagocytosis, cytokine responses, and immune activation (Figure 5I). While the large number of intergenic regions open in obese group relative to leans at T3 (Figure 5D) failed to enrich to any particular GO process by GREAT, motif analysis suggests enhanced binding sites for factors that regulate regulatory M2-like macrophages (STAT6 and LXR) (Figure 5J).

### Pregravid obesity is associated with metabolic and functional rewiring of circulating monocytes at term

The initiation of proinflammatory effector mechanisms requires a shift of cellular metabolism towards aerobic glycolysis. Therefore, we asked if the reduced response to LPS is associated with a lack of metabolic plasticity. Baseline extracellular acidification rate (ECAR) measurements suggested reduced cellular preference for glycolysis with pregravid obesity (n=3/group) (Figure 6A). We also observed attenuated induction of ECAR following LPS stimulation and glucose injection (Figure 6B). Furthermore, monocytes from subjects with obesity accumulated higher levels of neutral lipids (Figure 6C).

**Figure 6:**
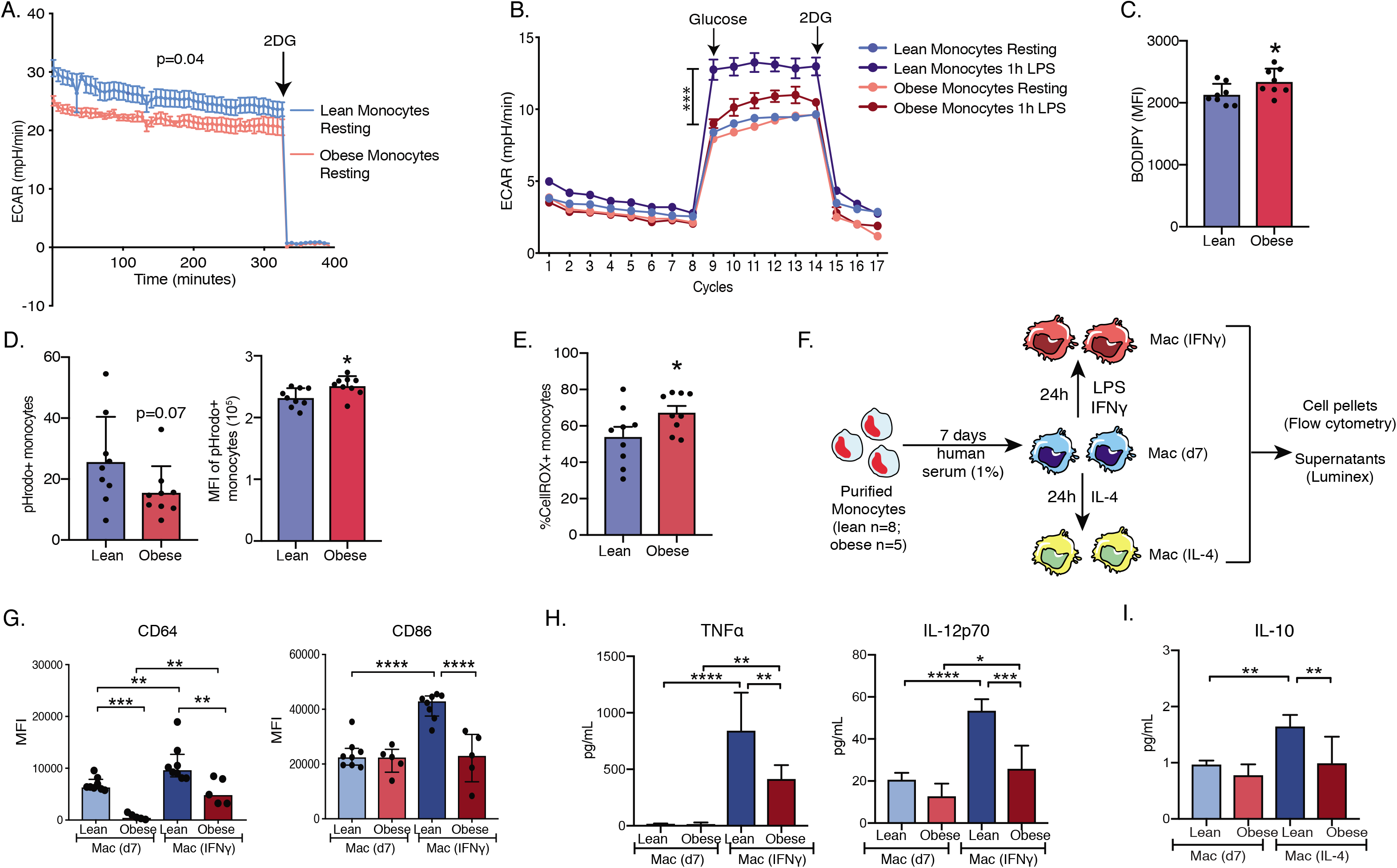
Metabolic and functional reprogramming of monocytes with maternal obesity at term. (A) Mean ECAR of purified monocytes from lean (blue) and obese (red) group (n=3/group) under basal conditions using glycolytic rate assay. (B) Mean ECAR of LPS activated monocytes (acute injection) from lean (blue) and obese (red) under glucose-free conditions, and post glucose injection using glycolytic stress assay. (C) Bar graphs comparing MFIs of BODIPY within CD14+ gate in PBMC (D) Bar graphs comparing pHrodo+ CD14+ cells following incubation of PBMC with pHrodo conjugated E.coli for 4 hours (left) and pHrodo signal within the pHrodo+ monocytes (right). (E) Cytosolic ROS readouts in LPS activated monocytes using FACS analysis of CellROX treated PBMC. (F) Experimental design for monocyte differentiation and polarization assay. (G) Bar graphs comparing induction of surface CD64 and CD86 expression following LPS and IFNγ stimulation of monocyte derived macrophages on day 7 post differentiation. (H) Secreted TNFα and IL-12p70 levels following LPS and IFNγ stimulation on day 7 using luminex. (I) Secreted IL-10 levels following IL-4 stimulation on day 7 post differentiation.

Additionally, we detected changes in effector functions in monocytes with pregravid obesity. Fewer cells from the obese group phagocytosed E.coli (Figure 6D, left), albeit the number of pathogen particles/cell was significantly higher (Figure 6D, right). Furthermore, pregravid obesity resulted in higher levels of cytosolic ROS following LPS stimulation (Figure 6F and Supp Figure 8A).

### Maternal obesity alters the differentiation trajectory of circulating monocytes at T3

Monocytes under state of immune tolerance (reduced responsiveness to LPS challenge) or recovering from tolerance (O’Carroll et al., 2014) demonstrate defects in differentiation and polarization trajectory. Therefore, we next investigated the impact of pregravid obesity on macrophage polarization potential using *in vitro* differentiation of monocytes collected at T3 using IFNγ (M1 like) and IL-4 (M2 like) conditioning (Figure 7F). Monocytes from mothers with obesity showed dampened induction of surface CD64 (Figure 7G, left) and CD86 (Figure 7G, right) following IFNγ stimulation with no differences in HLA-DR induction (Supp Figure 8B). In line with attenuated M1 skewing, secretion of pro-inflammatory cytokines TNFα and IL-12p70 (Figure 7H) and chemokines CXCL9 (Supp Figure 8C) and CXCL11 (Supp Figure 8D) was reduced following IFNγ stimulation. Although we observed no differences in the expression of M2-like macrophage surface markers CD163 (Supp Figure 8E) and CD209 (Supp Figure 8F) following IL-4 conditioning, we observed lower production of M2-like macrophage immune molecules IL-10 (Figure 7E), CCL11 (Supp Figure 8G) and PDGF-BB (Supp Figure 8H).

**Figure 7:**
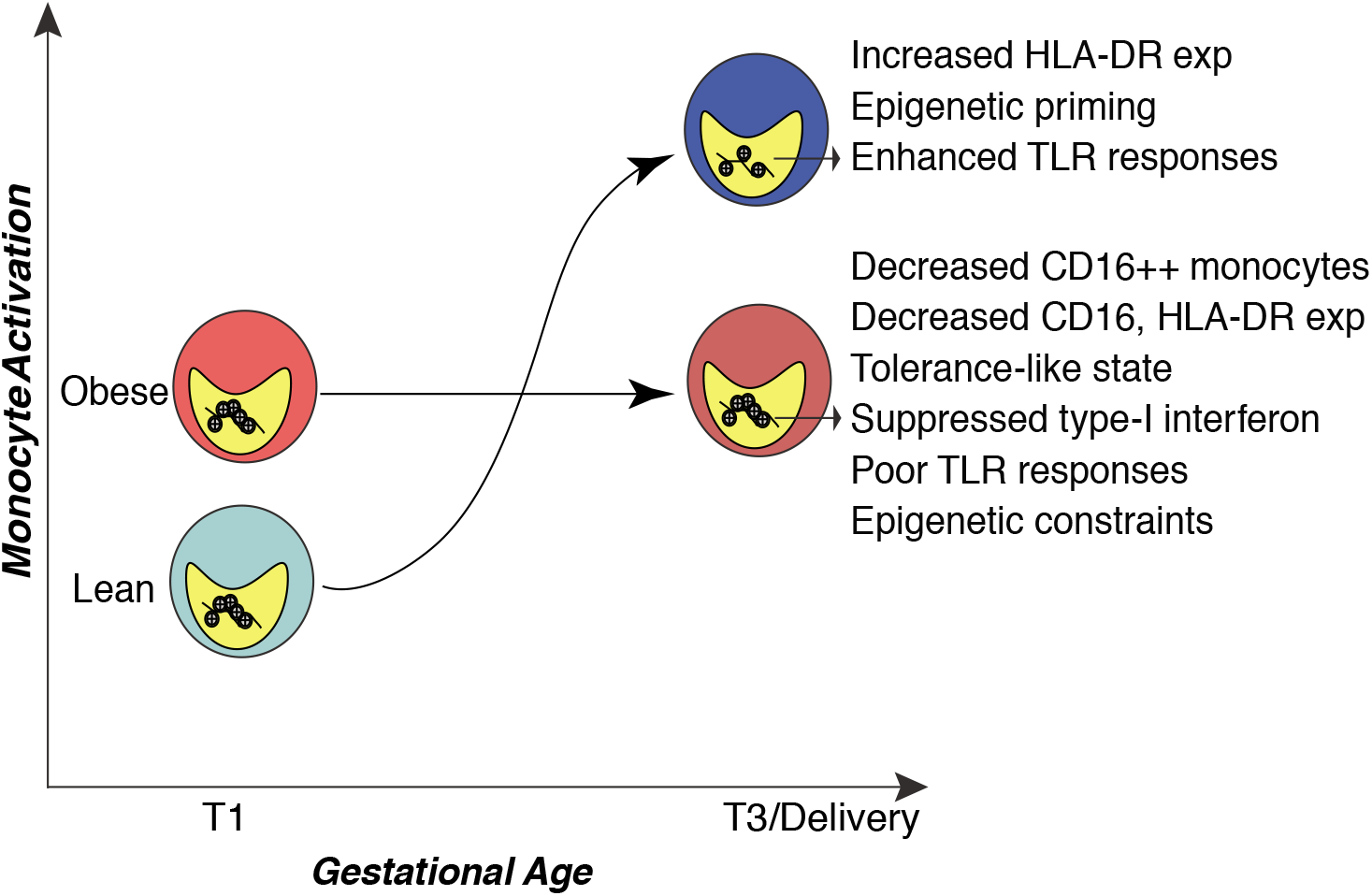
Model of the trajectory of monocyte activation with gestation and its disruption by pregravid obesity.

## DISCUSSION

Maternal obesity poses significant risks to maternal health, including increased susceptibility to infections during pregnancy (Robinson et al., 2005; Sebire et al., 2001; Stapleton et al., 2005), labor (Acosta et al., 2012; McLean et al., 2012; Salim et al., 2012) and post-partum (Acosta et al., 2012; McLean et al., 2012; Salim et al., 2012), suggesting disruptions in the pregnancy “immune clock”. This phenomenon refers to precisely timed adaptations in the maternal immune system, notably increased neutrophil numbers, enhanced innate immune responses, augmented STAT signaling in CD4+ T cells and NK cells, and significant changes in the circulating proteome (Aghaeepour et al., 2017; Aghaeepour et al., 2018; Le Gars et al., 2016). Previous studies profiling the impact of pregravid obesity on maternal peripheral immune have focused on one particular type of measurement at one stage of pregnancy (Acosta et al., 2012; McLean et al., 2012; Sureshchandra et al., 2018). Therefore, in this study, we used a systems approach to characterize the impact of pregravid obesity on pregnancy associated “immunological clock”, using a combination of proteomic, functional, and genomic assays.

Profiling longitudinal differences in plasma inflammatory factors between the first (T1) and third (T3) trimesters in lean subjects and those with pregravid obesity (BMI > 30) showed that Independent of maternal BMI, the transition from T1 to T3 was associated with increased levels of several pro-inflammatory (TNFα, S100B, IL-2, IFNβ, IL-17A, IL-18, IL-23) and regulatory (IL-4, IL-10, and IL-13) factors. These observations are consistent with previous reports of systemic immune activation and counter-regulation over the course of pregnancy (Aghaeepour et al., 2017). In addition to these analytes, and as previously reported (Basu et al., 2011a; Catalano et al., 2009; Challier et al., 2008; Friis et al., 2013; Roberts et al., 2011; Sen et al., 2014; Stewart et al., 2007), pregravid obesity resulted in significantly higher levels of IL-6 and CRP both at T1 and T3, highlighting a state of chronic low-grade inflammation. Higher levels of IL-6 and CRP have been linked to obstetric complications (Christian and Porter, 2014; Stewart et al., 2007).

Plasma levels of insulin, leptin, and adiponectin also increased with gestational age in lean subjects, likely a consequence of metabolic adaptations to the nutritional demands of a growing fetus. In contrast, in subjects with obesity, systemic levels of insulin, leptin, and resistin did not increase with gestational age. Aberrant levels of insulin and leptin have been associated with both miscarriage (Baban et al., 2010) and preeclampsia (Laivuori et al., 2000), both of which are increased with pregravid obesity.

Profiling circulating immune cell subsets at T1 and T3 revealed modest cellular adaptations. Pregnancy was associated with increased total WBC counts, as previously reported (Chandra et al., 2012). However, at both T1 and T3, mothers with obesity had higher WBC counts, relative to lean mothers. This significant increase was contributed by higher granulocyte counts, as previously described for non-gravid obese individuals (Rosales, 2018). No differences in T or B cell frequencies were observed with obesity. The most significant change with pregnancy was an increase in CD56^++^CD16^+^ subset of NK cells. This subset of regulatory NK cells has been shown to increase with gestational age, and is the dominant NK cell subset (>95%) in the placental decidua (Le Bouteiller, 2013). As reported in previous studies (Aghaeepour et al., 2017), no significant differences were seen in other innate immune cell subsets.

Interestingly, levels of plasma regulatory factors (IL1RA, PDGF, CCL22) were significantly elevated whereas those of IL-27, which activates monocytes via STAT1 signaling pathway, were lower in plasma of mothers with obesity at T3. Moreover, circulating levels of pro-inflammatory IL-1β increased with gestation in lean mothers but remained unchanged in mothers with obesity. Additionally, levels of CCL2 and CRP decreased with gestational age in mothers with obesity. Collectively, these observations strongly suggest that pregravid obesity disrupts changes in immune factors associated with myeloid cell activation, potentially to protect the developing fetus from obesity-induced chronic low-grade inflammation.

Indeed, pregravid obesity was associated with a lower frequency of nonclassical (CD16+) monocytes and lower expression of HLA-DR on monocytes at T3. Comparison of monocyte transcriptomes at T3 from lean mothers and those with obesity using high resolution single cell RNA sequencing (scRNA-Seq) confirmed a significant reduction in the number of non-classical monocytes, and expression of both *CD16* and *HLA-DR.* Additionally, scRNA-Seq analysis showed increased expression of genes encoding inflammatory molecules *LYZ*, *VCAN* and *HLA-DR* in monocytes from lean subjects, while expression of antiviral genes was suppressed in monocytes from obese subjects. Additionally, gene encoding regulatory molecules such as *CD163, AHR,* and *CD52* was increased with pregravid obesity.

These transcriptional changes were mirrored by functional differences in monocytes obtained from lean subjects and those with obesity at T3. Specifically, while activation circulating monocytes in lean subjects increased between T1 and T3 as indicated by increased frequency of IL-6/TNFα producing cells following LPS stimulation and elevated levels of plasma sCD14. Moreover, bulk RNA-Seq revealed an enhanced transcriptional response to LPS at T3 relative to T1 in the lean subjects relative to those with obesity marked by higher expression of pro-inflammatory molecules (*IL6*, *IL1B, TNF*) and induction of transcription factors (RELA, IRF1, STAT3). These observations are in line with earlier studies that have demonstrated that monocyte responses to influenza are enhanced with gestational age (Le Gars et al., 2016). Previous studies have reported increased endogenous STAT5ab signaling in T cells with pregnancy that support the development of regulatory T cells (Aghaeepour et al., 2017). Thus, enhancement of innate immune responses over the course of pregnancy might be a mechanism to counter dampened T cell responses and maintain anti-microbial immunity in the pregnant woman.

Pregravid obesity disrupted pregnancy-associated monocyte activation, with a significantly reduced number of TNFα/IL-6 producing monocytes relative to lean subjects in response to LPS at T3. Similarly, RNA Seq analysis revealed dampened induction of cytokine and chemokine genes as well as failure to upregulate expression of key transcription factors NFKB1 and FOSB following LPS stimulation. The dampened monocyte response is further evident from reduced surface expression of activation markers CD40 and CD86. Collectively, these differences in the innate immune system provide a potential explanation for the increased incidence of viral infections in obese pregnant women. It is possible that the state of chronic low-grade inflammation associated with obesity in combination with pregnancy-induced increase in circulating immune mediators result in a state of immune tolerance in circulating monocytes.

Several factors have been proposed to be responsible for monocyte activation with pregnancy including elevated levels of leptin and other cytokines (Sacks et al., 2001), exposure to placental microparticles released into circulation by syncytial trophoblasts (Redman et al., 2012), fetal DNA (Bianchi et al., 1996), or circulation of peripheral monocytes through the placenta (Mellembakken et al., 2002). Our RNA seq analysis also indicated changes in the expression of various epigenetic modifying factors. Supporting this state of enhanced activation with pregnancy, we report a significant increase in chromatin accessibility within promoter and intergenic loci that regulate inflammatory responses between T1 and T3 in resting monocytes from lean subjects. Epigenetic changes with healthy pregnancy have been shown in human uterine NK cells (Gamliel et al., 2018) and mouse mammary glands (Dos Santos et al., 2015). Our study is the first to highlight chromatin remodeling in circulating immune cells, providing the first clues of an epigenetic clock of human pregnancy. While the precise mechanisms regulating this epigenetic clock are not clear, studies have demonstrated that both type I interferons and TNF can cooperatively reprogram the epigenome of myeloid cells resulting in increased chromatin accessibility at inflammatory loci (Park et al., 2017) and establishment of transcriptional memory at the chromatin level. Indeed, our analysis of circulating immune mediators shows dramatic increase in plasma levels of both TNF and type-I interferons with pregnancy.

Pregravid obesity altered this epigenetic clock, resulting in a failure in the induction of transcripts associated with chromatin remodeling following LPS stimulation at T3, suggesting potential epigenetic constraints. Indeed, minimal chromatin changes were noted in monocytes from subjects with obesity between T1 and T3. A direct comparison of profiles at T1 and T3 shows a lack of remodeling within of promoter and enhancer regions that regulate cytokinesignaling (*TNF),* myeloid cell activation (*CD80*), and immune effector process (*HLA-DRA*) in monocytes from subjects with obesity. Moreover, regions that were accessible in monocytes from lean subjects (but remained closed in obese subjects) contained putative binding sites for transcription factors that orchestrate pro-inflammatory responses (AP-1, IRF1, IRF3, and IRF8). In contrast, regions that were accessible in monocytes from subjects with obesity harbored binding sites for regulatory factors such as STAT6 and SMAD2.

Epigenetic priming mechanisms play critical roles in tolerance and training of innate immune memory, and our findings support the hypothesis that pregravid obesity results in tolerance in circulating monocytes. Specifically, expression of *HLA-DR* in monocytes at T3 measured at the protein, transcript, and epigenetic levels were significantly reduced. Lower MHC class II molecule expression has been described in models of endotoxin tolerance and in sepsis patients during late stages of immunoparalysis (Wolk et al., 2003).

An immunotolerant state has been shown to affect functional plasticity in monocytes. Earlier studies have reported enhanced intracellular ROS levels (Sacks et al., 1998) but reduced phagocytic function (Lampe et al., 2015) in monocytes with pregnancy. Functional assessment of purified monocytes at T3 showed that pregravid obesity results in a reduction in the number of monocytes that can phagocytose bacteria but an increased aggregate pathogen engulfment per cell. Furthermore, monocytes from pregnant subjects with obesity had an enhanced oxidative burst following LPS exposure. This is in contrast to studies describing reduced *ex vivo E. coli* and *S. aureus* induced ROS production in tolerant monocytes (Grondman et al., 2019). These differences in outcomes could be explained by different triggers of oxidative stress (LPS vs. pathogen). Immune tolerance can also impact the ability of monocytes to differentiate towards M1- or M2-like state under different polarization conditions. In agreement with transcriptional and cytokine responses, monocyte-derived macrophages (MoDM) from mothers with obesity polarized poorly to M1 but not to M2 skewing conditions. Cytokine secretion, however, were dampened under both conditions, suggesting overall reduced plasticity compared to the lean counterparts.

Another manifestation of immune tolerance is the lack of metabolic plasticity. Indeed, monocytes obtained from humans injected with low dose LPS show a lack of metabolic plasticity, failing to up-regulate glycolysis, pentose phosphate pathway (PPP), and down-regulate lipid metabolic pathways when rechallenged *ex vivo* with LPS (Grondman et al., 2019). Our analysis of maternal monocytes at T3 showed reduced basal extracellular acidification rate (ECAR) with obesity indicative of dampened preference for glycolysis. Additionally, we observed poor induction of glycolysis following LPS injection. This coupled with the accumulation of higher levels of neutral lipids further supporting metabolic reprogramming with maternal obesity.

In summary, this study revealed that pregravid obesity disrupts the monocyte activation trajectory associated with pregnancy (Figure 7), thereby providing a potential explanation for increased susceptibility to infections during gestation and post-partum in pregnant subjects with obesity. The study also provides a conceptual framework of an epigenetic clock that supports the immune clock of pregnancy. We were able to demonstrate that pregravid obesity disrupts the pregnancy immune clock, altering the metabolic, molecular and functional phenotype of peripheral monocytes towards a regulatory phenotype. Future studies will focus on generating a comprehensive model of monocyte activation with additional gestational time points as well as assessing the impact of pregravid obesity on the monocyte response to pathogens. Peripheral blood monocytes are recruited to become decidual macrophages; hence future experiments will also interrogate the impact of pregravid obesity at the maternal fetal interface.

## MATERIALS AND METHODS

### Subjects and experimental design

This study was approved by the Institutional Ethics Review Board of Oregon Health and Science University and the University of California, Irvine. Written consent was obtained from all subjects. A total of 117 non-smoking women (69 lean and 48 obese) who had an uncomplicated, singleton pregnancy were enrolled for this study. The breakdown is as follows: 1) 37 women classified as lean and 28 women classified as obese were enrolled during first trimester (~week 12, designated as T1) of whom 29 lean and 27 obese also provided a blood sample during the third trimester (~Week 37, designated as T3); 2) 32 women classified as lean and 20 women classified as obese based on prepregnancy BMI were enrolled during the third trimester and provided one sample at ~37 weeks (T3). The mean age of the 69 lean women was 32.8±4.7 years and the average pre-pregnancy BMI was 21.9 ± 1.7 kg/m^2^; mean age of the 48 obese women was 29.4±9 years with an average pre-pregnancy BMI of 35.2 ± 5.2 kg/m^2^ (Table 1).

The racial distribution of the entire cohort was as follows: 101 Caucasian, 7 Asian American, 2 Pacific Islander, 6 American Indian/Alaskan native, 6 African American, and 3 declined to report. Exclusion criteria included active maternal infection, documented fetal congenital anomalies, substance abuse, chronic illness requiring regular medication use, preeclampsia, gestational diabetes, chorioamnionitis, significant medical conditions (active cancers, cardiac, renal, hepatic, or pulmonary diseases), or an abnormal glucose tolerance test. Women underwent a fasting blood draw and body composition via air displacement plethysmography using a BodPod (Life Measurement Inc).

### Plasma and Peripheral Blood Mononuclear Cell (PBMC) isolation

Complete blood counts were obtained by Beckman Coulter Hematology analyzer (Brea, CA). Peripheral blood mononuclear cells (PBMC) and plasma were obtained by standard density gradient centrifugation over Ficoll (GE Healthcare). PBMC were frozen in 10% DMSO/FBS and stored in liquid nitrogen until analysis. Plasma was stored at −80°C until analysis.

### Luminex assay

Immune mediators in plasma were measured using a customized multiplex human factor panel (R & D Systems, Minneapolis MN) measuring cytokines (IFNβ, IFNγ, IL-1β, IL-10, IL-12p70, IL-13, IL-15, IL-17A, IL-18, IL-1RA, IL-2, IL-21, IL-4, IL-5, IL-7, TNFα, IL-23, IL-31, IL-22, IL-27), chemokines (CCL2/MCP-1, CCL3/MIP-1α, CCL4/MIP-1β, CCL5/RANTES, CCL11/Eotaxin, CXCL1/GROα, CXCL8/IL-8, CXCL9/MIG CXCL10/IP-10, CXCL11/I-TAC, CXCL12/SDF-1α, CXCL13/BCA-1, growth factors (BDNF, GM-CSF, HGF, EGF, VEGF, PDGF-BB) and additional molecules (PD-L1, S100). Metabolic hormones were measured using a 3-plex kit measuring insulin, leptin, and PYY (Millipore, Burlington MA). Adipokines were assayed using a 5-plex kit measuring adiponectin, adpisin, lipocalin-2, total PAI-1, and resistin (Millipore, Burlington MA). Samples were diluted per manufacturer’s instructions and run in duplicates on the Magpix Instrument (Luminex, Austin, TX). Data were fit using a 5P-logistic regression on xPONENT software.

### ELISA

CRP and IL-6 were measured using a high sensitivity ELISA (Life Technologies, Carlsbad CA). Plasma soluble CD14 (sCD14) was measured using ELISA per manufacturer’s instruction (Hycult Biotech, Uden, Netherlands)

### PBMC and monocyte phenotyping

10^6^ PBMC were stained using the following cocktail of antibodies to enumerate innate immune cells and their subsets: PE-CD3, PE-CD19, PB-CD16, PE-Cy7-CD11c, AF700-CD14, PCP-Cy5.5-CD123, BV711-CD56, and APC-Cy7-HLA-DR. An additional 10^6^ PBMC from a subset of samples were stained using an additional panel to further characterize monocyte phenotype: AF700-CD14, APC-Cy7-HLA-DR, PB-CCR5, AF488-TLR2, BV711-TLR4, PCP-Cy5.5-CD163, BV605-CCR2, PE-eF610-CD11c, PE-Cy7-CD11b, and PE-CX3CR1. After surface staining, cell pellets were washed twice in PBS and resuspended in cold FACS buffer (PBS with 2% FBS and 1mM EDTA). All samples were acquired with the Attune NxT Flow Cytometer (ThermoFisher Scientific, Waltham MA) and analyzed using FlowJo 10.5.0 (Ashland OR).

### Intracellular cytokine assay and monocyte activation

To measure cytokine responses of monocytes and dendritic cells, 10^6^ PBMC were stimulated for 16h at 37°C in RPMI supplemented with 10% FBS in the presence or absence of 1 ug/mL LPS (TLR4 ligand, *E.coli* 055:B5; Invivogen, San Diego CA); Brefeldin A (Sigma, St. Louis MO) was added after 1 hour incubation. Cells were stained for APC-Cy7-CD14 and PCP-Cy5.5-HLA-DR, fixed, permeabilized, and stained intracellularly for APC-TNF *α* and PE-IL-6. To measure immune activation, a subset of PBMC samples were stimulated with LPS for 16h without Brefeldin A, washed twice with PBS and surface stained using a cocktail of following antibodies for 20 minutes: FITC-CD62L, AF700-CD14, HLA-DR-APC-Cy7, PB-CD16, BV510-CD40, PE-Cy7-CD80, PCP-Cy5.5-CD83, BV605-CD86, BV711-CD64, PE-CXCR6, and PE-Dazzle594-PD-L1. Cell pellets were washed twice in PBS and resuspended in cold FACS buffer (PBS with 2% FBS and 1mM EDTA). All samples were acquired with the Attune NxT Flow Cytometer (ThermoFisher Scientific, Waltham MA) and analyzed using FlowJo 10.5 (Ashland OR).

### Isolation of monocytes

Monocytes were purified from freshly thawed PBMC using CD14 antibodies conjugated to magnetic microbeads per the manufacturer’s recommendations (Miltenyi Biotec, San Diego CA). Magnetically bound monocytes were washed and eluted for collection. Purity was assessed using flow cytometry and was on average ?95% (Supp Figure 7A).

### Monocyte stimulation

10^5^ purified monocytes were cultured in RPMI supplemented with 10% FBS with or without 1 ug/mL LPS (TLR4 ligand, *E.coli* 055:B5; Invivogen, San Diego CA) in 96-well tissue culture plates at 37C in a 5% CO_2_ environment for 8 hours. Plates were spun down and cell pellets were resuspended in Qiazol (Qiagen) for RNA extraction. Both cells and supernatants were stored in −80°C until they could be processed as a batch.

### Library generation for bulk RNA-Seq

Total RNA was isolated from monocytes using mRNeasy kit (Qiagen, Valencia CA). Quality and concentrations were measured using Agilent 2100 Bioanalyzer. Libraries were generated using TruSeq Stranded Total RNA-Seq kit (Illumina, San Diego CA). Briefly, following rRNA depletion, mRNA was fragmented for 8 min, converted to double stranded cDNA and adapter ligated. Fragments were then enriched by PCR amplification and purified. Size and quality of the library was verified using Qubit and Bioanalyzer. Libraries were multiplexed and sequenced on the HiSeq4000 platform (Illumina, San Diego CA) to yield an average of 20 million 100 bp single end reads per sample.

### Bulk RNA-Seq analysis

Quality control of raw reads was performed retaining bases with quality scores of ≥20 and reads ≥35 base pair long. Reads were aligned to human genome (hg38) using splice aware aligner TopHat using annotations available from ENSEMBL (GRCh38.85) database. Quantification of read counts was performed using GenomicRanges package in R and normalized to derive transcripts per million (TPM) counts. To detect the effect of pregravid obesity in resting monocytes at T1 and T3 respectively, differential expression analysis was performed using quasi-likelihood linear modeling in edgeR. To test pairwise longitudinal changes with lean and obese groups, data was fit into negative binomial GLMs with a design matrix that preserved sample ID information. Lowly expressed genes were filtered at the count stage, eliminating ones with 0 counts in at least 3 samples regardless of the group. Due to the relatively low number of genes passing the standard FDR cutoff (FDR <0.05), genes with expression difference (relaxed statistical p-value <0.01) were included in subsequent analyses.

Responses to LPS were modeled pairwise at each time point using negative binomial GLMs following low read count filtering. Genes with expression difference (FDR <0.05) were considered differentially expressed genes (DEG). Functional enrichment of DEG was performed using Metascape and InnateDB. Transcription factors that regulate expression of DEG were predicted using ChEA3 tool using ENSEMBL ChIP database. Heatmaps of fold change or TPMs were generated and bubble plots for enrichment of Transcription Factor (TF) regulation or Gene Ontology (GO) terms was generated using ggplot in R. Gene expression data have been deposited in NCBI’s Sequence Read Archive (SRA accession number pending).

### Cell Sorting and library generation for single cell (sc)RNA-seq

PBMC from delivery time point were thawed and live cells were sorted into RPMI (supplemented with 30% FBS) using the BD FACS Aria Fusion and SYTOX Blue (1:1000, ThermoFisher). Sorted cells were counted in triplicates and resuspended in PBS with 0.04% BSA in a final concentration of 1200 cells/uL. Cells were immediately loaded in the 10X Genomics Chromium with a loading target of 17,600 cells. Libraries were generated using the V3 chemistry per the manufacturer’s instructions (10X Genomics, Pleasanton CA). Libraries were sequenced on Illumina NovaSeq with a sequencing target of 50,000 reads per cell.

### scRNA-seq data analysis

Raw reads were aligned and quantified using the Cell Ranger Single-Cell Software Suite (version 3.0.1, 10X Genomics) against the GRCh38 human reference genome using the STAR aligner. Downstream processing of aligned reads was performed using Seurat (version 3.1.1). Droplets with ambient RNA (cells fewer than 400 detected genes), potential doublets (cells with more than 4000 detected genes, and dying cells (cells with more than 20% total mitochondrial gene expression) were excluded during initial QC. Data objects from lean and obese group were integrated using Seurat. Data normalization and variance stabilization was performed using *SCTransform* function using a regularized negative binomial regression, correcting for differential effects of mitochondrial and ribosomal gene expression levels and cell cycle. Dimension reduction was performed using *RunPCA* function to obtain the first 30 principal components followed by clustering using the *FindClusters* function in Seurat. Visualization of clusters was performed using UMAP algorithm as implemented by Seurat’s *runUMAP* function. Cell types were assigned to individual cluster using *FindMarkers* function with a fold change cutoff of at least 0.5 and using a known catalog of well-characterized scRNA markers for PBMC (Zheng et al., 2017).

Three monocyte clusters expressing high levels of *CD14* were extracted from dimension reduced Seurat object using the *subset* function. Cells were reclustered and visualized using UMAP. Doublet clusters were identified by iterative clustering removing clusters with high expression of genes associated with T cells, B cells, and NK cells. Differential expression analysis was tested using Wilcoxon rank sum test followed by bonferroni correction using all genes in the dataset. For gene scoring analysis, we compared gene signatures and pathways from KEGG (https://www.genome.jp/kegg/pathway.html) in subpopulations using Seurat’s *AddModuleScore* function. Two-way functional enrichment of differential signatures was performed on Metascape. Differential hierarchies within the monocyte compartment were reconstructed using Monocle (version 2.8.0). Briefly, clustering was performed using t-SNE and differential genes identified using Monocle’s *differentialGeneTest.* These genes (q-value < 1e-10) were used for ordering cells on a pseudotime.

### Neutral Lipid Staining

Neutral lipids in monocytes were quantified in monocytes using flow cytometry. Briefly, 500,000 PBMC were surface stained (CD14-AF700, HLA-DR-APC-Cy7) for 20 minutes at 4C, washed twice, and resuspended in 500 uL warm 1X PBS containing 1 ug/mL BODIPY™ 493/503 (ThermoFisher Scientific). Cells were incubated at 37C for 10 minutes and samples were acquired with the Attune NxT Flow Cytometer (ThermoFisher Scientific, Waltham MA) and analyzed using FlowJo 10.5 (Ashland OR).

### Metabolic Assays

Basal Oxygen Consumption Rate (OCR) and Extracellular Acidification Rate (ECAR) were measured using Seahorse XF Glycolysis Rate Assay on Seahorse XFp Flux Analyzer (Agilent Technologies) following manufacturer’s instructions. Briefly 200,000 purified monocytes were seeded on Cell-Tak (Corning) coated 8-well culture plates in phenol free RPMI media containing 2 mM L-glutamine, 10 mM L-glucose, 1mM sodium pyruvate, and 5mM HEPES. Seeded plates were placed in 37C incubator without CO_2_ and run on the XFp with extended basal measurements, followed by serial injections of Rotenone/Actinomycin (final well concentration 0.5 uM) to block oxidative phosphorylation followed by 2-Deoxy-D-Glucose (2-DG, 500 mM) to block glycolysis. Cellular responses to stress under activated conditions were measured using Seahorse XF Glycolysis Stress Assay. Purified monocytes were seeded in the in glucose free media and cultured in the presence/absence of 1 ug/mL LPS for 1 hour in 37C incubator without CO_2_. Plates were run on the XFp for 8 cycles of basal measurements, followed by an acute injection of L-glucose (100 mM), oligomycin (50 uM), and 2-DG (500 mM). Analysis and interpretation of data was done on Seahorse Wave desktop software (Agilent Technologies).

### Phagocytosis Assay

Cellular phagocytosis was measured using pH-sensitive pHrodo® E. coli BioParticles® conjugates (ThermoFisher Scientific). 500,000 PBMC were activated with 100 ng/mL LPS for 4h, washed twice and incubated for an additional 2 hours in media containing pHrodo conjugates at 1 mg/mL conjugates. Pellets were washed twice, surface stained (CD14-AF700, HLA-DR-APC-Cy7), and resuspended in ice-cold FACS buffer. All samples were acquired with the Attune NxT Flow Cytometer (ThermoFisher Scientific, Waltham MA) and analyzed using FlowJo 10.5 (Ashland OR).

### Cellular ROS assay

For intracellular ROS measurements, 500,000 PBMC were stimulated with 1 ug/mL LPS for 4 hours. For negative control, cells were incubated in serum-free media containing 200 uM anti-oxidant N-acetylcysteine (NAC) for 1.5 hours. Both negative and positive controls were incubated with tert-butyl hydroperoxide (TBHP) for 30 minutes to induce oxidative stress. All samples were then incubated with 2.5 uM CellROX Deep Red (Life Technology) at 37C for 30 minutes, washed twice, surface stained (CD14-FITC, HLA-DR-PCP-Cy5.5) and samples were acquired with the Attune NxT Flow Cytometer (ThermoFisher Scientific, Waltham MA) and analyzed using FlowJo 10.5 (Ashland OR).

### *In vitro* Macrophage Differentiation

100,000 purified monocytes were cultured in 24-well tissue culture plates treated for increased attachment (VWR) in RPMI supplemented with 1% Human AB Serum (Omega Scientific) for 7 days with media supplemented on day 3. On day 7, cells were polarized to M1-like macrophages using 1 ug/mL LPS and 100 ng/mL IFNγ (Peprotech) or M2-like macrophages using 100 ng/mL IL-4 (Peprotech) and cultured for an additional 24 hours (day 8). Cell pellets from day 7 and day 8 were surface stained (CD16-PB; CD64-BV711; HLA-DR-APC-Cy7; CD86-BV605; CD163-PCP-Cy5.5; CD209-R-PE) and analyzed using flow cytometry. Supernatants were collected and quantified for secreted cytokines and chemokines using a 29-plex luminex assay (R & D Systems).

### Library generation for ATAC-Seq

ATAC-Seq libraries were generated using a recently described modified protocol (OMNI-ATAC) to reduce mitochondrial reads. Briefly, 50,000 purified monocytes were lysed in lysis buffer (10mM Tris-HCl (pH 7.4), 10 mM NaCl, 3 mM MgCl_2_), for 3 min on ice to prepare the nuclei. Immediately after lysis, nuclei were spun at 500g for 10 min to remove the supernatant. Nuclei were then incubated with transposition mixture (100 nM Tn5 transposase, 0.1% Tween-20, 0.01% Digitonin and TD Buffer) at 37C for 30 min. Transposed DNA was then purified with AMPure XP beads (Beckman Coulter) and partially amplified for 5 cycles using the following PCR conditions – 72C for 3 min; 98C for 30s and thermocycling at 98C for 10s, 63C for 30s and 72C for 1 min. To avoid overamplification, qPCR was performed on 5 uL of partially amplified library. Additional cycles of amplification for the remainder of the sample were calculated from the saturation curves (cycles corresponding to a third of the saturation value). Fully amplified samples were purified with AMPure beads and quantified on the Bioanalyzer (Agilent Technologies, Santa Clara CA).

### Analysis of ATAC-Seq data

Paired reads from sequencing were quality checked using FASTQC and trimmed to retain reads with quality scores of ≥20 and minimum read lengths of 50 bp. Trimmed paired reads were aligned to the human genome (hg38) using Bowtie2 (–X 2000 –k 1 –very-sensitive –no-discordant –no-mixed). Reads aligning to mitochondrial genome and allosomes were removed using samtools. PCR duplicate artifacts were then removed using Picard. Finally, poor quality alignments and improperly mapped reads were filtered using samtools (samtools view –q 10 –F 3844). To reflect the accurate read start site due to Tn5 insertion, BAM files were repositioned using ATACseqQC package in R. The positive and negative strands were offset by +4bp and −5bp respectively. Samples within a group were merged and sorted using samtools.

Sample QC and statistics for merged BAM files were generated using HOMER makeTagDirectory function. Accessible chromatin peaks were called for mapped paired reads using HOMER findpeak function (-minDist 150 -region —fdr 0.05). PCA and sample clustering were performed on accessible peaks using DiffBind. Differentially accessible regions (DAR) in either direction were captured using HOMER getDiffererentialPeaks function (-q 0.05). DAR were annotated using the human GTF annotation file (GRCh38.85) and ChIPSeeker with a promoter definition of −1000 bp and +100 bp around the transcriptional start site (TSS). Peaks overlapping 5’UTRs, promoters, first exons and first introns were pooled for functional enrichment of genes. For intergenic changes, the genes closest to the intergenic DAR were considered. Functional enrichment of this pooled list of genes was performed using DAVID (Fisher p-value <0.05). BAM files were converted to bigWig files using bedtools and visualized on the new WashU EpiGenome browser.

### Statistical analysis

All statistical analyses were conducted in Prism 8 (GraphPad). All definitive outliers in two-way and four-way comparisons were identified using ROUT analysis (Q=0.1%). Data was then tested for normality using Shapiro-Wilk test (alpha=0.05). If data were normally distributed across all groups, differences with obesity and pregnancy were tested using ordinary one-way ANOVA with unmatched samples. Multiple comparisons were corrected using Holm-Sidak test adjusting the family-wise significance and confidence level at 0.05. If gaussian assumption was not satisfied, differences were tested using Kruskall-Wallis test (alpha=0.05) followed by Dunn’s multiple hypothesis correction test. Differences in normally distributed datasets were tested using an unpaired t-test with Welch’s correction (assuming different standard deviations). Two group comparisons that failed normality tests were carried out using Mann-Whitney test. Associations and correlograms between BMI and cytokine levels were generated using corrplot package in R.

## Acknowledgements

We thank Samantha Castañeda and Andrew N. Tang for assistance with immune assays; Selene B. Nguyen for assistance with preparation of RNA-Seq libraries; and Brian Jin Kee Ligh for help with ATAC-Seq data analysis. We thank Dr. Jennifer Atwood for assistance with sorting and imaging flow cytometry in the flow cytometry core at the Institute for Immunology, UCI. We thank Dr. Melanie Oakes from UCI Genomics and High-Throughput Facility for assistance with 10X library preparation and sequencing.

## Author Contributions

SS, NEM, and IM conceived and designed the experiments. SS, NM, AJ, and MZ performed the experiments. SS, IM and NM analyzed the data. SS, NEM and IM wrote the paper.

## Declaration of Interests

The authors have declared that no financial or non-financial competing interests exist.

## Funding

This study was supported by grants from the National Institutes of Health 1K23HD06952 (NEM), R03AI112808 (IM), 1R01AI142841 (IM), and 1R01AI145910 (IM).

## Data availability

The datasets supporting the conclusions of this article are available on NCBI’s Sequence Read Archive (SRA# pending).

## SUPPLEMENT FIGURES

**Supplementary Figure 1: Longitudinal changes in circulating inflammatory environment.**

(A) Linear regression of fat mass and pregravid BMI at T1 and T3. (B) Dot plots demonstrating gestational weight gain (GWG) in lean and obese subjects. (C-F) Bar graphs comparing circulating levels of (C) resistin, (D) adipsin, (E) adiponectin, and (F) peptide YY (PYY) at T1 and T3. (G-I) Four-way violin plots comparing plasma levels of (G) IL-27, (H) CCL4, and (I) VEGF-A. (J) Association between changes in plasma analyte level with gestation (T3-T1) and BMI. Size of the bubble reflects the strength of Pearson’s correlation, and color indicates direction of association (red-positive; blue-negative). Levels of significance: * – p<0.05, ** – p<0.01; *** – p<0.001; ****– p<0.0001.

**Supplementary Figure 2: Immune cell profiling**

(A) Complete blood counts of all blood samples analyzed (B) Gating strategy for characterization of innate immune cell populations from PBMC. (C-E) Bar graphs comparing frequencies of total (C) monocytes, (D) dendritic cells, and (E) NK cells within PBMC. (F-G) Comparing frequencies of (F) myeloid DCs subsets, (G) plasmacytoid DCs, and (H) NK cell subsets. Levels of significance: *–p<0.05, ** – p<0.01.

**Supplementary Figure 3: Phenotyping of adaptive immune cells** (A-C) Percentages of total (A) B cells, (B) CD4+ T cells, and (C) CD8+ T cells within PBMC. (D-F) Relative abundances of naïve and memory subpopulations with (D) B cells, (E) CD4+ T cells, and (F) CD8+ T cells. Levels of significance: *– p<0.05, ** – p<0.01, **** – p<0.0001.

**Supplementary Figure 4: Cytokine responses to *ex vivo* stimulation.** (A) Gating strategy for measuring frequencies of responding monocytes and T cells following ex vivo stimulation using intracellular cytokine staining. (B-D) Violin plots of responding T cells following CD3/CD28 stimulation. (B) IL-2 (above) and IL-4 (below) responses in CD4+ T cells. IFNγ (above) and TNFα (below) responses in (C) CD4+ and (D) CD8+ T cells.

**Supplementary Figure 5: Baseline transcriptional differences with pregnancy.** (A-B) Scatter plot depiction of normalized transcript counts (TPM) on a log2 scale from resting monocyte at T1 and T3 (projected on X and Y axes respectively) obtained from (A) lean and (B) obese groups. Only genes with significant expression differences (p<0.001) are annotated. (C) UMAP visualization of PBMC isolated from lean mothers (n=2) and mothers with obesity (n=2) at delivery time point. Live cells were enriched using FACS and profiled using droplet based single cell RNA sequencing. (D) Feature plots of characteristic markers to delineate monocyte subsets based on their levels of expression as gradient of purple. (E) Pseudotime ordering of monocytes (left) revealing progressive shifts in cellular states with pregravid obesity (right). (F-G) Violin plots comparing expression of genes involved in (F) type-I interferon signaling across different clusters, highlighting select (G) interferon stimulated genes (ISGs) significantly down-regulated with maternal obesity.

**Supplementary Figure 6: Profiling transcriptional responses to LPS.**

(A) Gating strategy for assessment of purity of monocytes following magnetic bead separation. (B) Comparing LPS responsive DEG in lean and obese group at T1 and (C) T3.

**Supplementary Figure 7: Epigenetic adaptations with pregnancy and obesity**

(A) Functional enrichment of genes regulated by promoter associated and (B) intergenic DAR open in lean group at T3 relative to T1 (FC>2). Genes associated with these regions are quantified next to each GO term. (C) Genomic contexts of DAR comparing monocyte ATAC peaks from leans and obese group at T1. (D-E) WashU Epigenome tracks for (D) *IL6ST* and (E) *CD80* locus with promoter vicinity highlighted in yellow. (F) Gene ontologies of intergenic associations identified by GREAT as significantly open in obese group relative to lean group at T3. (G) WashU Epigenome tracks for *EHMT1* significantly more open with obesity.

**Supplementary Figure 8: Functional rewiring of monocytes with maternal obesity at term**

(A) Contour plots of CellROX signal from monocytes gated in lean and obese samples. NAC and TBHP treated monocytes serve as negative and positive controls respectively. (B) Bar graphs comparing surface expression of HLA-DR and secreted levels of M1 associated chemokines (C) CXCL9 and (D) CXCL11 following LPS and IFNγ stimulation on day 7. (E) Bar graphs comparing surface expression of M2 associated markers CD163 and (F) CD209, and M2-associated chemokine (G) CCL11 (eotaxin) and growth factor (H) PDGF. Levels of significance: * – p<0.05; *** – p<0.001; ****-p<0.0001. Bars represent medians and interquartile ranges.

